# The effects of aging on neural signatures of temporal regularity processing in sounds

**DOI:** 10.1101/522375

**Authors:** Björn Herrmann, Chad Buckland, Ingrid S. Johnsrude

## Abstract

Sensitivity to temporal regularity (e.g., amplitude modulation) is crucial for speech perception. Degradation of the auditory periphery due to aging and hearing loss may lead to an increased response gain in auditory cortex, with potential consequences for the processing of temporal regularities. We used electroencephalography recorded from younger (18–33 years) and older (55–80 years) adults to investigate how aging affects neural gain and the neural sensitivity to amplitude modulation in sounds. Aging was associated with reduced adaptation in auditory cortex, suggesting an age-related gain increase. Consistently, neural synchronization in auditory cortex to a 4-Hz amplitude modulation of a narrow-band noise was enhanced in ~30% of older individuals. Despite enhanced responsivity in auditory cortex, sustained neural activity (likely involving auditory and higher-order brain regions) in response to amplitude modulation was absent in older people. Hence, aging may lead to an over-responsivity to amplitude modulation in auditory cortex, but to a diminished regularity representation in higher-order areas.

## Introduction

When more than one sound source is present at a time in a listener’s environment, the sounds mix and the auditory system faces the challenge of decomposing this mixture into meaningful parts (e.g., a person’s voice or an approaching car) so that one particular sound source can be identified and tracked. Each individual sound has a unique profile of amplitude modulations. In speech, for example, low-frequency (<10 Hz) amplitude modulations reflect the unique evolution of the word/syllable envelope of an utterance (Rosen, 1992). Tracking such modulations is thought to support speech comprehension in the presence of other sounds (Kerlin et al., 2010; Giraud and Poeppel, 2012; Peelle and Davis, 2013). Understanding how amplitude modulations in sounds are processed is thus essential for understanding success and failure during listening to real-world auditory signals including speech.

Hearing impairment is common among normally aging adults (Cruickshanks et al., 1998; Feder et al., 2015; Goman and Lin, 2016). Older people have difficulty hearing very quiet sounds (due to elevated hearing thresholds) (Moore, 2007; Plack, 2014), but they also experience difficulty with suprathreshold sounds and this is more relevant for their everyday function. Issues include comprehension difficulties when speech is heard in the presence of background sound (Helfer and Wilber, 1990; Pichora-Fuller and Souza, 2003), perception of sounds at moderate intensities to be unpleasantly loud (Epstein and Marozeau, 2006; Tyler et al., 2014), and increased distraction by irrelevant sounds (Parmentier and Andrés, 2010; Mishra et al., 2014). These issues may be due, at least in part, to altered salience of amplitude modulation in sound. In fact, older people with hearing impairment have better detection thresholds for faint low-frequency (<10 Hz) amplitude modulations in suprathreshold sounds (Ernst and Moore, 2012; Schlittenlacher and Moore, 2016) and may perceive suprathreshold amplitude modulations to fluctuate more strongly compared to normal-hearing individuals (Moore et al., 1996).

Work in animals suggests that hearing loss and aging is accompanied by changes throughout the auditory system. Accumulated noise exposure is thought to damage the synapses that connect inner hair cells with auditory nerve fibers (Kujawa and Liberman, 2009; Bharadwaj et al., 2014; Viana et al., 2015; Liberman and Kujawa, 2017), which in turn reduces the input from the auditory periphery to neural circuits in brainstem, thalamus, and cortex. A consequence of peripheral damage is a loss of neural inhibition throughout the auditory system (including cortex) (Caspary et al., 2008; Llano et al., 2012; Takesian et al., 2012; Auerbach et al., 2014). Consistent with a loss of inhibition following peripheral damage, neural responses to sounds are enhanced in the aged and noise-exposed auditory system of animals (Hughes et al., 2010; Chambers et al., 2016; Salvi et al., 2017) and humans (Laffont et al., 1989; Herrmann et al., 2013b; Bidelman et al., 2014; Herrmann et al., 2016; Henry et al., 2017). A loss of inhibition is hypothesized to compensate for reduced inputs from the periphery by up-regulating the responsivity to sound in subcortical and cortical circuits (Caspary et al., 2008; Takesian et al., 2012); this is referred to as central gain enhancement/increase (Auerbach et al., 2014).

In human listeners, changes in inhibition and central gain may be assessed by measuring sound-evoked neural adaptation (Herrmann et al., 2016; Herrmann et al., 2018). Neural adaptation refers to the reduction of a neural response caused by repetitive sound stimulation (Malmierca et al., 2014; Nelken, 2014). How neurons recover from adaptation can be measured by varying the time interval between two successive stimulus presentations, where a longer inter-stimulus interval is associated with a longer recovery time and thus a larger response to a subsequent stimulus (Hari et al., 1982; Sams et al., 1993; Zacharias et al., 2012; Herrmann et al., 2016). Critically, neural responses increase more strongly with longer inter-stimulus intervals in older compared to younger human adults, suggesting that neurons in the aging auditory cortex recover faster from adaptation (Herrmann et al., 2016). This reduction in neural adaptation is thought to be a consequence of reduced cortical inhibition (Herrmann et al., 2016; Herrmann et al., 2018).

How a loss of inhibition and increased responsivity affects the processing of low-frequency temporal regularities such as amplitude modulations in sounds is less well understood. Sensitivity to temporal regularity in sounds can be assessed by the strength of neural synchronization (Purcell et al., 2004; Herrmann et al., 2013a; Goossens et al., 2016; Henry et al., 2017). Neural synchronization reflects the alignment of neural activity with periodicity in sound (Lakatos et al., 2008; Stefanics et al., 2010; Lakatos et al., 2013; Henry and Herrmann, 2014; ten Oever et al., 2017), and is strongest in auditory cortex for low-frequency (<10 Hz) periodicities (Herrmann et al., 2013a; Keitel et al., 2017; Millman et al., 2017). Some studies report increased synchronization of neural activity with low-frequency amplitude modulations in sounds for older compared to younger humans (Purcell et al., 2004; Goossens et al., 2016; Presacco et al., 2016a) and non-human mammals (Overton and Recanzone, 2016; Herrmann et al., 2017; Lai et al., 2017; Parthasarathy et al., 2019). This enhanced synchronization may reflect a loss of inhibition.

Sensitivity to temporal regularity may also be assessed by the magnitude of sustained neural activity (Barascud et al., 2016; Sohoglu and Chait, 2016; Teki et al., 2016; Southwell et al., 2017; Herrmann and Johnsrude, 2018a). Sustained neural activity is a low-frequency DC power offset that occurs when a listener hears a regular pattern in a sound, such as repeating sequences of tones with random frequencies or trains of isochronous clicks (Gutschalk et al., 2002; Barascud et al., 2016; Sohoglu and Chait, 2016; Teki et al., 2016; Southwell et al., 2017; Herrmann and Johnsrude, 2018a). Sustained activity and neural synchronization are dissociable, since they are differently susceptible to the attentional state of listeners (Herrmann and Johnsrude, 2018a). Neural synchronization decreases, whereas sustained neural activity increases, when listeners attend to sounds. Moreover, neural synchronization and sustained neural activity may also involve distinct neural generators. Neural synchronization originates from auditory cortex (Herrmann et al., 2013a; Keitel et al., 2017), whereas sustained activity is thought to originate from higher-level brain regions including frontal cortex, parietal cortex, and hippocampus (Tiitinen et al., 2012; Barascud et al., 2016; Teki et al., 2016) in addition to auditory cortex (Pantev et al., 1994; Pantev et al., 1996; Gutschalk et al., 2002). Increases in sustained activity may reflect a prediction-related increase in sensitivity when uncertainty in sound environments is reduced (Gutschalk et al., 2002; Barascud et al., 2016; Heilbron and Chait, 2018). However, it is unclear how aging affects regularity-related sustained neural activity and the relative magnitudes of synchronized and sustained activity.

The current study investigates whether the neural signatures of temporal regularities in sounds differ between younger and older people. By utilizing a sound that contains a temporally regular amplitude modulation, the effects of age on sensory representation (indexed by neural synchronization), as well as on regularity representations that may involve higher-level cortices (indexed by sustained activity) can be examined concomitantly (Herrmann and Johnsrude, 2018a). Age-related changes in neural processing are investigated in three ways: (1) We utilize a stimulus protocol that assesses the temporal dynamics of neural adaptation in auditory cortex, since reduced inhibition in older people may manifest as faster adaptation recovery times in auditory cortex (Herrmann et al., 2016). (2) We assess the tendency for neural activity to synchronize with a sound’s amplitude modulation, since reduced inhibition in aging also may also be reflected in enhanced neural synchronization to low-frequency amplitude modulation in sounds (Purcell et al., 2004; Goossens et al., 2016; Presacco et al., 2016a, b; Herrmann et al., 2017). (3) Neural processing that may involve additional higher-level brain regions is assessed using the sustained neural activity in response to the same amplitude-modulated sounds that we utilize to assess sensory gain enhancements. We hypothesize that if reduced inhibition and enhanced sensory gain degrades neural representations of a sound’s regularity, then sustained neural activity should be reduced, despite enhanced sensory representation (i.e., reduced neural adaptation and enhanced neural synchronization).

## Methods and Materials

### Participants

30 younger (mean: 23.7 years, range: 19–33 years, 18 female) and 26 older (mean: 67.1 years, range: 55–76 years, 16 female) adults participated in the current study. Four additional participants were excluded: two because of extensive movement during the EEG recordings; one because of a Montreal Cognitive Assessment (MoCA; Nasreddine et al., 2005) screening score that indicated mild cognitive impairment (<26), and which may affect neural activity in sensory areas (Bidelman et al., 2017); and one because of severe hearing impairment as identified using audiometry (see below). Participants reported no neurological disease or hearing impairment and were naïve to the purposes of the experiment. Participants gave written informed consent prior to the experiment and were paid $5 CAD per half-hour for their participation. The study was conducted in accordance with the Declaration of Helsinki, the Canadian Tri-Council Policy Statement on Ethical Conduct for Research Involving Humans (TCPS2-2014), and was approved by the local Nonmedical Research Ethics Board of the University of Western Ontario (protocol ID: 106570).

### Stimulation apparatus

All experimental procedures were carried out in a sound-attenuating booth. Sounds were presented via Sennheiser (HD 25-SP II) headphones and a Steinberg UR22 (Steinberg Media Technologies) external sound card. Stimulation was controlled by a PC (Windows 7, 64 bit) running Psychtoolbox in Matlab (R2015b).

### Hearing assessment

Pure-tone audiometry was assessed for each participant for six frequencies: 0.25, 0.5, 1, 2, 4, and 8 kHz. Participants with a low-frequency hearing loss greater than 30 dB HL (average across 0.25, 0.5, 1, 2 kHz frequencies) were excluded from the current study (N=1).

Participants filled out a detailed questionnaire to assess their subjective experience related to speech perception and loudness perception. We used a questionnaire made publically available in a recent study (Liberman et al., 2016). The questionnaire assessed self-rated speech perception in quiet (one question) and noise (four questions), and spatial hearing (three questions) (similar to SSQ questions; Gatehouse and Noble, 2004). Participants rated these questions on a 10-point scale (with 10 indicating ‘perfect’ agreement). We also assessed the participants’ subjective loudness experience, sound annoyance experience, and sound avoidance behavior. These questions were rated on a 100-point scale (with 100 indicating ‘unbearable loud’, ‘annoying’, and ‘avoiding’, respectively). The full questionnaire can be downloaded at https://osf.io/7snrz/.

### Sensation level and loudness perception

Before the experiment, the sensation level (i.e., hearing threshold) was determined for each participant for a 1272-Hz sine tone using a method-of-limits procedure. The threshold was used as a reference point to assess each participant’s range for loudness perception. Thresholds were obtained by presenting sounds of 15-s duration that changed continuously in intensity at 4.33 dB/s (either decreased [i.e., starting at suprathreshold levels] or increased [i.e., staring at subthreshold levels]). Participants indicated via button press when they could no longer hear the tone (intensity decrease) or when they started to hear the tone (intensity increase). The mean sound intensity at the time of the button press was noted for 6 decreasing sounds and 6 increasing sounds (decreasing and increasing sounds alternated), and these were averaged to determine the individual hearing threshold.

Peripheral (cochlea) function was assessed by measuring loudness perception in a category scaling experiment (Oxenham and Bacon, 2003; Epstein and Marozeau, 2006; Al-Salim et al., 2010; Hebert et al., 2013; Plack, 2014). Participants listened to tones, one at a time, and indicated the perceived loudness of that tone using one of seven categories (‘not heard’, ‘very soft’, ‘soft’, ‘comfortable’, ‘loud’, ‘very loud’, or ‘too loud’). Participants were instructed to judge the loudness of each tone subjectively, based on that presentation alone, and not to compare it with preceding tone presentations. Tones were of 0.7 s duration were presented at one of 22 different sound levels ranging from −1.67 dB to 80 dB (step size: 3.89) relative to the hearing threshold measured from a young, normal-hearing participant group (N>100) in a previous study (Herrmann and Johnsrude, 2018b). Hearing thresholds of normal-hearing ears relate to about 20 dB SPL (Moore et al., 1996). Each sound level was repeated 6 times and presented randomly. In order to quantify the range across which sound levels were perceived, we calculated the loudness range as (1) the difference in the level (in dB) for tones perceived as ‘very soft’ relative to ‘very loud’, and (2) the difference in the level between a participant’s hearing threshold (from the psychophysical method-of-limits procedure) and the level for tones perceived as ‘very loud’. We did not use the ‘too loud’ category for the calculation of the loudness range because not all participants used this category. An independent samples t-test compared the loudness range between age groups.

### Acoustic stimulation for electrophysiological measurement of temporal adaptation dynamics

In order to measure the temporal dynamics of neural adaptation, we closely followed the experimental design from our previous study (Herrmann et al., 2016). To this end, sequences of 0.1-second pure tones (at 1272 Hz; 7 ms linear rise and 7 ms linear fall time) were presented, each 18 tones long. The onset-to-onset interval changed logarithmically, becoming progressively shorter (from 3.8 s to 0.4 s) in one half of the sequence and then progressively longer in the other half (to 2.959 s; see Figure 1A). Twenty sequences were concatenated to make one long sequence. The duration of the onset-to-onset interval preceding a tone served as the independent variable by which trials were sorted into conditions. Presentation of the concatenated sequence was repeated three times in separate blocks (each approx. 9 min) and participants listened passively while watching a muted movie of their choice with subtitles on a battery-driven portable DVD player. Over the experiment, 1080 tones were presented (18 tones × 20 sequences = 360 × 3 repetitions = 1080). For two older participants, no data were recorded for this part of the study, because the participants felt uncomfortable sitting for the extended period required by the experimental procedures. As a consequence, only 24 data sets from older people were available for data analysis of the temporal dynamics of adaptation.

**Figure 1:**
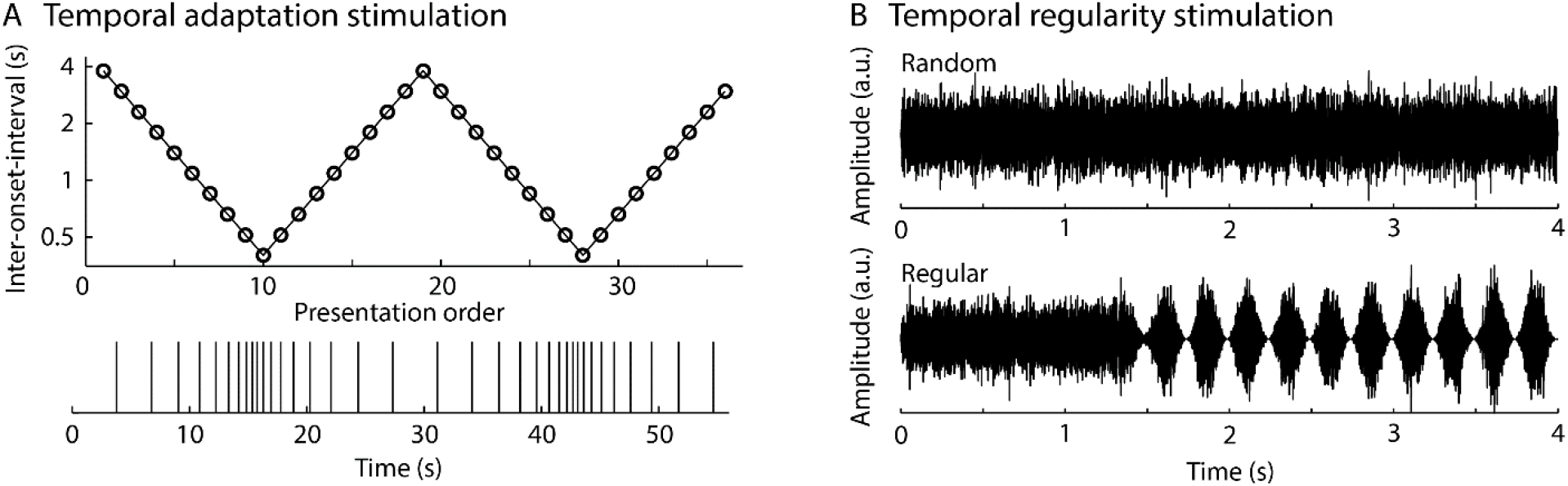
Experimental stimulation to measure temporal dynamics of neural adaptation (A) and temporal regularity processing (B).

### Acoustic stimulation for electrophysiological measurement of temporal regularity processing

Stimuli were narrow-band noises made by adding 50 amplitude-modulated pure-tone components of 4-second duration. For each of the 50 components, a random carrier frequency between 900 Hz and 1800 Hz was selected (starting phase was random). The amplitude of each component was modulated at a randomly changing rate between 2 Hz and 6 Hz (average rate of 4 Hz; different components were modulated at different rates). The 50 components were summed in order to obtain a narrow-band noise. We henceforth refer to this stimulus as the ‘random’ condition (see Figure 1B, top). A ‘regular’ condition was created by manipulating the number of sound components with consistent phase. In this condition, the amplitude modulation of all 50 components aligned in phase at about 1.55 seconds after sound onset, and the amplitude modulation rate remained constant at 4 Hz for the rest of the sound. Stimuli were created such that there was no discontinuity at the transition from non-aligned to phase-aligned parts of the sound. The waveform of a ‘regular’ example is displayed in Figure 1B (bottom).

Participants listened passively to 150 trials of each condition (random, regular) while watching a muted movie of their choice with subtitles on a battery-driven portable DVD player. The experiment was divided into 3 blocks (each approx. 9 min long), separated by short breaks during which participants rested. During each block, 50 trials of each condition were randomly presented. Trials were separated by a 1.5 s inter-stimulus interval.

### Intensity of sounds during EEG recording

Sounds played in the two EEG experiments (one measuring temporal adaptation dynamics, and one measuring temporal regularity processing; the order was counterbalanced across participants) were presented at 60 dB above the average hearing threshold for a group of young, normal-hearing adults (N>100) (Herrmann and Johnsrude, 2018b). For 10 out of the 56 participants, sounds were presented at 60 dB relative to their individual hearing threshold. We quantified the intensity of the sounds played during EEG recordings relative to each individual’s hearing threshold (i.e., sensation level) and relative to each individual’s comfortable listening level, which was derived from the loudness category scaling experiment.

Relative to sensation level, sounds played to participants during the EEG recording were on average louder for younger compared to older participants (t_53_ = 4.097, p < 0.001, r_e_ = 0.490; Figure 3B). Furthermore, the intensity of sounds played during the EEG recording was slightly louder relative to the comfortable sound level for younger participants (t_29_ = 3.147, p = 0.004, r_e_ = 0.505), but not different from the comfortable sound level for older participants (t_25_ = 1.384, p = 0.179, r_e_ = 0.267; the difference between age groups was not significant: t_54_ = 1.569, p = 0.122, r_e_ = 0.209; Figure 3B). The fact, that the sounds presented during the EEG recordings were, on average, louder for younger compared to older participants works against our hypothesis that sensory gain is enhanced in older people. Sound level is thus not a confounding factor in our study.

### EEG recordings and preprocessing

EEG signals were recorded at a 1024-Hz sampling rate from 16 electrodes (Ag/Ag–Cl-electrodes; 10-20 placement) and from the left and right mastoids (BioSemi, Amsterdam, The Netherlands; 208 Hz low-pass filter). Electrodes were referenced to a monopolar reference feedback loop connecting a driven passive sensor and a common mode sense active sensor, both located posteriorly on the scalp.

Offline data analysis was carried out using MATLAB software (v7.14; MathWorks, Inc.). Data from one electrode (‘O2’) were excluded for all participants, because the electrode broke over the course of the study. Line noise (60 Hz) was suppressed in the raw data using an elliptic filter. Data were re-referenced to the average mastoids, high-pass filtered at a cutoff of 0.7 Hz (2449 points, Hann window), and low-pass filtered at a cutoff of 30 Hz (111 points, Hann window). Independent components analysis (ICA; runica method, Makeig et al., 1996; logistic infomax algorithm, Bell and Sejnowski, 1995; Fieldtrip implementation Oostenveld et al., 2011) was used to identify and suppress activity related to blinks. Data recorded in temporal adaptation blocks (Figure 1A) were divided into epochs ranging from −0.05 to 0.3 s (time-locked to sound onset). Data recorded in temporal regularity blocks (Figure 1B) were divided into epochs ranging from −0.6 to 4.6 s (time-locked to sound onset). Epochs that exceeded a signal change of more than 200 μV for any electrode were excluded from analyses. This pipeline was used to investigate neural adaptation and neural synchronization (see below).

In order to investigate sustained neural activity, the same analysis pipeline was calculated a second time, with the exception that high-pass filtering was omitted. Omission of the high-pass filter is necessary to investigate the sustained response, which is a very low-frequency DC shift (Barascud et al., 2016; Southwell et al., 2017; Herrmann and Johnsrude, 2018a). Activity related to blinks was suppressed using the identified components from the high-pass filtered data.

Finally, the signal for the broken ‘O2’ electrode was interpolated by replacing it by the average signal across electrodes ‘Pz’, ‘P4’, and ‘O1’. The interpolated signal was used only for the display of topographical distributions.

### EEG data analysis: Temporal dynamics of neural adaptation

Response time courses for unique onset-to-onset intervals were obtained as follows. Single-trial time courses for a unique onset-to-onset interval and its direct neighbors (i.e., shorter and longer intervals) were binned and averaged (Ingham and McAlpine, 2005). For example, to obtain the average response for the 0.514-s interval, we averaged all trials for the 0.400-s, 0.514-s, and 0.660-s (n−1, n, n+1) onset-to-onset intervals. The overlap across intervals was used to increase the number of trials in the response average, thereby increasing the signal-to-noise ratio. This averaging procedure reduced the 10 logarithmically-spaced onset-to-onset intervals to 8 logarithmically-spaced onset-to-onset intervals (0.514, 0.660, 0.847, 1.088, 1.397, 1.794, 2.304, 2.959 s).

The analyses focused on N1 amplitudes for which neural adaptation has been previously reported (Hari et al., 1982; Sams et al., 1993; Zacharias et al., 2012; Herrmann et al., 2016). Mean amplitudes for a broad electrode cluster (Fz, F3, F4, Cz, C3, C4, Pz, P3, P4) were extracted for the N1 time window (0.8–0.11 s; Herrmann et al., 2016). In order to investigate the sensitivity of neural responses to interval duration, a linear function was used to relate N1 amplitudes to onset-to-onset intervals (separately for each participant). Onset-to-onset intervals were log-transformed before the linear function fit in order to account for the intervals’ logarithmic spacing (i.e., the log-transform linearized the interval spacing). The estimated linear coefficient from the linear fit reflects the sensitivity of the N1 amplitude to interval duration. Negative values index increasing N1 amplitudes with increasing interval duration. A larger negative coefficient for data from older compared to younger people would suggest faster recovery from adaptation over time in the auditory cortex of older people.

In order to assess whether N1 amplitudes were sensitive to interval duration, linear coefficients from the function fits were tested against zero using a one-sample t-test (separately for the younger and the older participant group). An independent samples t-test was used to compare linear coefficients between age groups.

### EEG data analysis: Neural synchronization to temporal regularity

Neural synchronization was operationalized as inter-trial phase coherence (ITPC; Lachaux et al., 1999). Single-trial time courses (high-pass filtered data) from the temporal regularity stimulation protocol (Figure 1B) were transformed into the frequency domain by calculating a fast Fourier transform (including a Hann window taper and zero-padding) for each trial, condition, and channel for the time window during which the temporal regularity (i.e., amplitude modulation) could occur in sounds: 1.55–4 s. The resulting complex numbers were normalized by dividing each complex number by its magnitude. ITPC was then calculated as the absolute value of the mean normalized complex number across trials. ITPC values can take on values between 0 (no coherence) and 1 (maximum coherence).

ITPC was calculated for frequencies ranging from 1–11 Hz (step size: 0.01 Hz). For statistical analyses, ITPC values were averaged across the fronto-central-parietal electrode cluster (Fz, F3, F4, Cz, C3, C4, Pz, P3, P4; Herrmann and Johnsrude, 2018a). ITPC values at the 4-Hz stimulation frequency were extracted by calculating the average across the 3.95–4.05 Hz frequency window. An analysis of variance (ANOVA) was carried out in order to compare neural synchronization strength (ITPC) between conditions (with-subject factor: random vs. regular) and age groups (between-subject factor: younger vs. older).

### EEG data analysis: Sustained neural activity

Single-trial time courses (non-high-pass filtered data) were averaged for each condition (random, regular) from the temporal regularity stimulation protocol (Figure 1B). Response time courses were baseline corrected by subtracting the mean amplitude in the pre-stimulus time window (−0.6 to 0 seconds) from the amplitude at each time point. In order to investigate the sustained response, signals were averaged across the fronto-central-parietal electrode cluster (Fz, F3, F4, Cz, C3, C4, Pz, P3, P4; Herrmann and Johnsrude, 2018a) and, for each condition, the mean amplitude was calculated for the 1.55–4 s time window during which the temporal regularity could occur (Barascud et al., 2016; Teki et al., 2016; Herrmann and Johnsrude, 2018a). An ANOVA was carried out in order to compare the amplitude of the sustained response between conditions (with-subject factor: random vs. regular) and age groups (between-subject factor: younger vs. older).

### Effect size

Throughout the manuscript, effect sizes are provided as partial eta squared (η_p_^2^) for ANOVAs and r_e_ (r_equivalent_) (Rosenthal and Rubin, 2003) for t-tests. re is equivalent to a Pearson product-moment correlation for two continuous variables, to a point-biserial correlation for one continuous and one dichotomous variable, and to the square root of partial η^2^ for ANOVAs.

## Results

### Hearing assessment

Audiometric thresholds are depicted in Figure 2A. All participants included in this study had an average hearing threshold below or equal to 30 dB HL in a low frequency region (average across 0.25, 0.5, 1, 2 kHz frequencies) at which sounds were presented. Older participants exhibited moderate to severe hearing loss at frequencies equal to and above 4 kHz.

**Figure 2:**
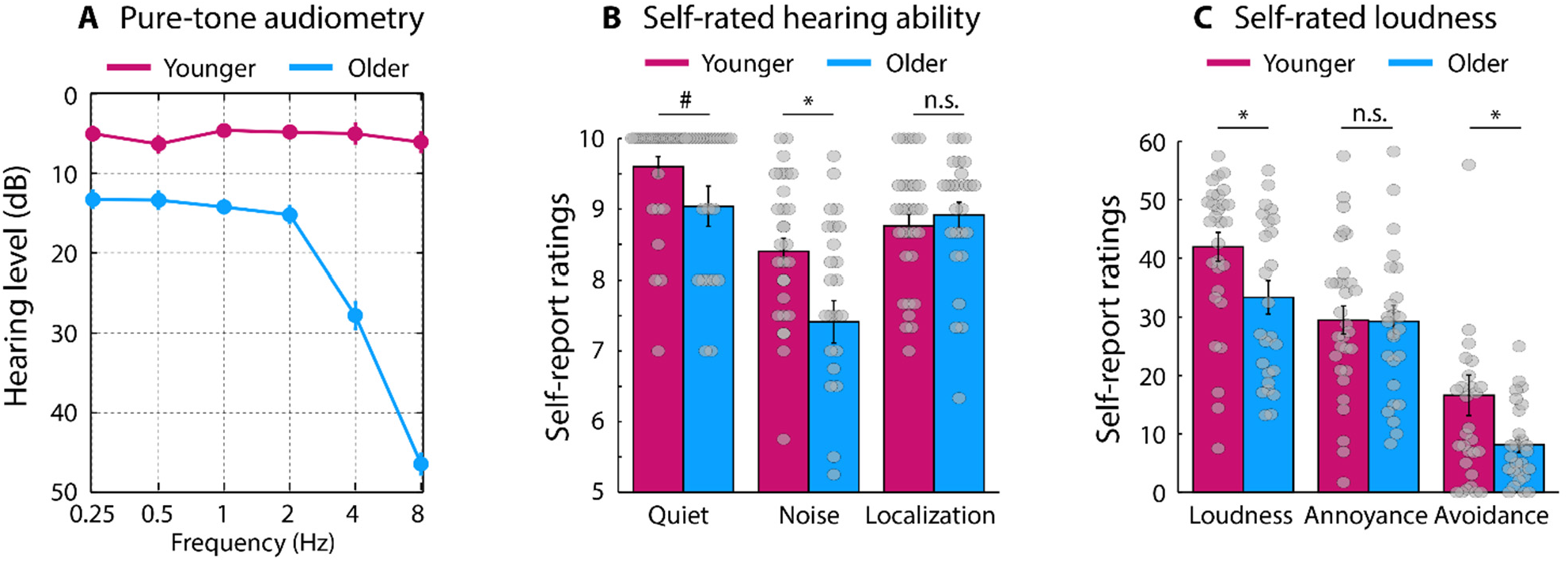
Hearing assessment measures. **A:** Audiometric thresholds. **B:** Self-rated hearing ability for speech in quiet, speech in noise, and spatial hearing. **C:** Self-rated loudness perception, sound annoyance, and avoidance of sound. Gray dots represent data points for individual participants. Error bars reflect the standard error of the mean. *p < 0.05, #p < 0.1, n.s. – not significant.

The results of a short version of the SSQ questionnaire (Gatehouse and Noble, 2004) assessing speech perception and spatial localization abilities are displayed in Figure 2B. Older people reported experiencing greater challenges when listening to speech in noisy environments than did younger people (t_54_ = 2.873, p = 0.006, r_e_ = 0.364; Figure 2B, middle). There was also a marginal effect of age for speech perception in quiet environments (t_54_ = 1.832, p = 0.073, r_e_ = 0.242; Figure 2B, left). No effect of age was found for self-rated spatial localization abilities (t_54_ = 0.648, p = 0.520, r_e_ = 0.088; Figure 2B, right).

The results from the questionnaire that assessed loudness perception, annoyance, and avoidance are depicted in Figure 2C. Unexpectedly, loudness ratings (t_54_ = 2.266, p = 0.028, r_e_ = 0.295) and avoidance ratings (t_54_ = 2.160, p = 0.035, r_e_ = 0.282) were lower for older compared to younger people. No age difference was found for annoyance ratings (t_54_ = 0.061, p = 0.952, r_e_ = 0.008). These results are in contrast to previous work using the same questionnaire for the assessment of loudness, annoyance, and avoidance in younger participant groups with and without a history of noise exposure (Liberman et al., 2016). Older people may interpret the situations described for the self-rating differently than do younger people.

### Loudness judgments

Loudness perception was assessed using a category scaling task (Epstein and Marozeau, 2006; Hebert et al., 2013). Participants categorized the loudness of tones into one of seven categories. Figure 3A shows the category judgment for each age group. Older participants perceived tones presented at low sound levels to be softer compared to younger participants. This is consistent and perhaps expected given the age-group difference in audiometric hearing thresholds for pure tones (Figure 2A). The difference in perceived loudness between age groups was not observed for more intense tones (Figure 3A). This is consistent with a reduced dynamic range for loudness in older compared to younger people (Hebert et al., 2013).

**Figure 3:**
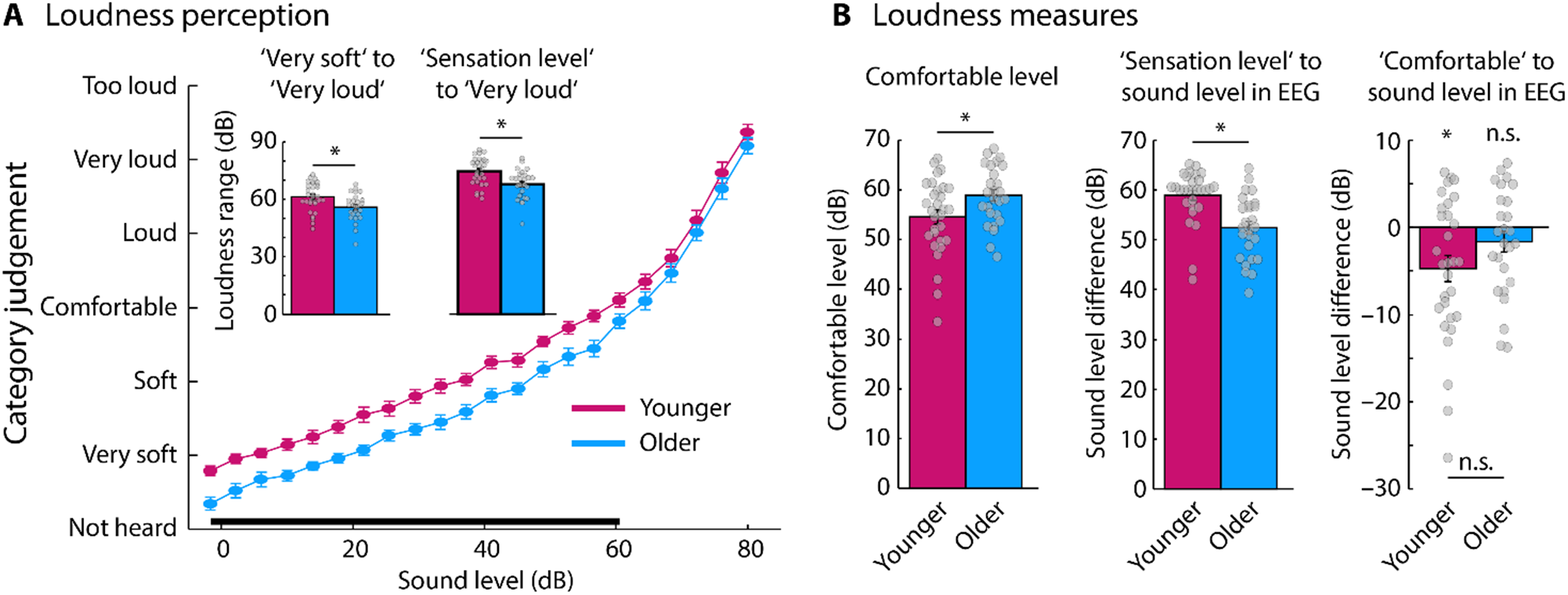
Results for loudness category judgments. **A:** Shows the mean loudness judgments for each age group (the black solid line indicates a significant age group difference, p < 0.05). The inset shows the two measures of loudness range (‘very soft’ to ‘very loud’ and ‘sensation level’ to ‘very loud’). Dynamic range for loudness was reduced in older compared to younger people. **B:** Different loudness measures calculated from the loudness judgments: comfortable level, level of sounds during EEG recordings relative to sensation level, and level of sounds during EEG recordings relative to comfortable level. Gray dots represent data points for individual participants. Error bars reflect the standard error of the mean. *p < 0.05, n.s. – not significant.

In order to test whether the loudness range differed between age groups, the perceptual loudness range was calculated. The loudness range for ‘very soft’ relative to ‘very loud’ percepts (t_54_ = 2.791, p = 0.007, r_e_ = 0.355), as well as for the hearing threshold relative to ‘very loud’ percepts (t_53_ = 3.361, p = 0.001, r_e_ = 0.419), was smaller in older compared to younger individuals (Figure 3A, inset). In addition, the comfortable sound level was more intense in older compared to younger people (difference: 4.33 dB; t_54_ = 2.328, p = 0.024, r_e_ = 0.302; Figure 3B, left).

### Neural adaptation

Figure 4A displays neural response time courses and topographical distributions for N1 responses (0.08–0.11 s time window). Topographical distributions are consistent with neural generators in auditory cortex (Näätänen and Picton, 1987; Picton et al., 2003). The degree to which neural responses recover from adaptation over time was investigated by fitting a linear function to N1 amplitudes as a function of the interval duration preceding a tone. The linear coefficient was significantly smaller than zero for younger (t_29_ = −9.752, p < 0.001, r_e_ = 0.875) and older people (t_23_ = −9.990, p < 0.001, r_e_ = 0.902), which shows that the N1 amplitude increases with increasing interval between two tones. The linear coefficient was larger (i.e., more negative) for older compared to younger participants (t_52_ = 2.252, p = 0.029, r_e_ = 0.298), which suggests that neurons in auditory cortex of older people recover faster from adaptation over time. There was no difference in overall amplitude between age groups (t_52_ = 1.049, p = 0.299, r_e_ = 0.144).

**Figure 4:**
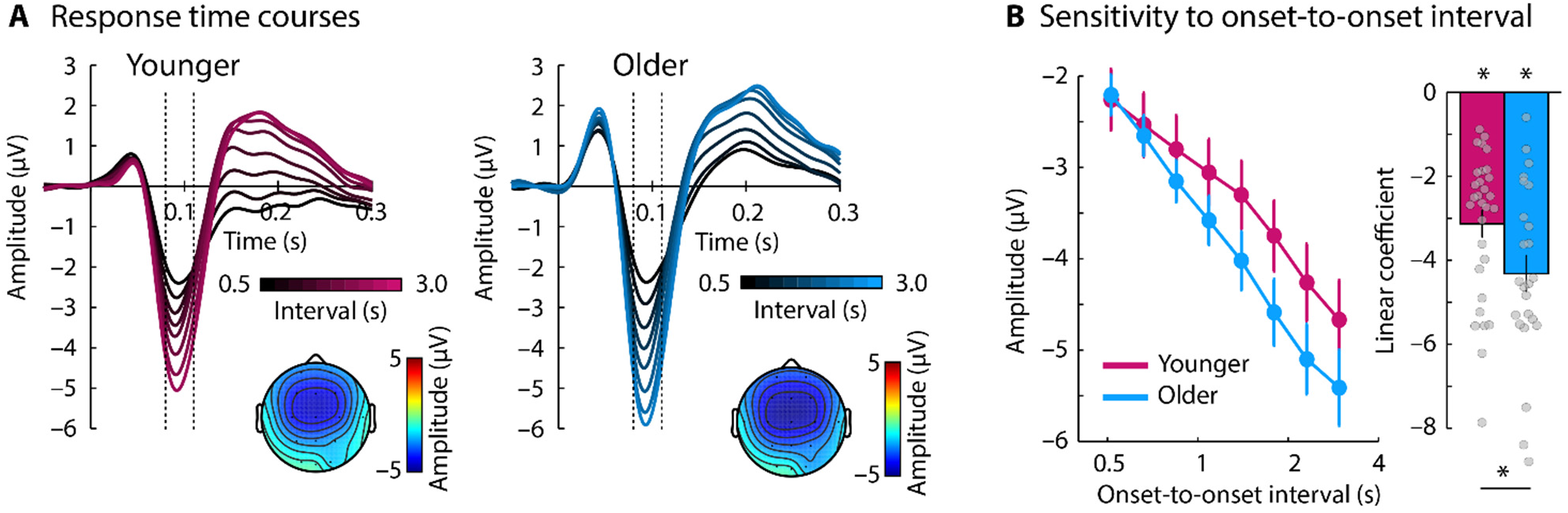
Results from the temporal adaptation design. **A:** Response time course for each onset-to-onset interval condition and age group. The vertical dashed lines mark the time interval of the N1 component (0.08–0.11 s). Topographies reflect the mean N1 response across interval conditions. **B:** Mean N1 response for each onset-to-onset interval condition and age group. The bar graphs on the right shows the mean linear coefficient (across participants) from the linear function fit to N1 amplitudes as a function of onset-to-onset interval. Gray dots represent the linear coefficients for individual participants. Error bars reflect the standard error of the mean. *p < 0.05.

### Neural synchronization to temporal regularity

Inter-trial phase coherence (ITPC) was calculated for stimuli that varied in phase coherence (temporal regularity stimuli). ITPC frequency spectra and topographies are depicted in Figure 5. ITPC at the 4-Hz stimulation frequency was compared between conditions and age groups. The ANOVA revealed a main effect of condition (F_1,54_ = 34.633, p < 0.001, η_p_^2^ = 0.391) and a main effect of age group (F_1,54_ = 5.707, p = 0.020, η_p_^2^ = 0.096), demonstrating larger ITPC for the ‘regular’ condition (i.e., sounds with coherent amplitude modulation) compared to the ‘random’ condition (i.e., sounds without coherent amplitude modulation), and larger overall ITPC for older people. Moreover, the condition × age group interaction was significant (F_1,54_ = 4.161, p = 0.046, η_p_^2^ = 0.072). ITPC was larger for the regular compared to the random condition for younger (t_29_ = 5.666, p < 0.001, r_e_ = 0.725) and older individuals (t_25_ = 3.956, p < 0.001, r_e_ = 0.621; Figure 5), but the difference in ITPC between the ‘regular’ and the ‘random’ condition was larger in older compared to younger individuals (t_54_ = 2.040, p = 0.046, r_e_ = 0.268). This is consistent with previous work showing an enhancement of neural synchronization for amplitude-modulated sounds at low modulation rates for older (>60 years) compared to younger people (<30 years) (Goossens et al., 2016; Presacco et al., 2016a, b). Topographical distributions are consistent with auditory cortex generators (Näätänen and Picton, 1987; Picton et al., 2003).

**Figure 5:**
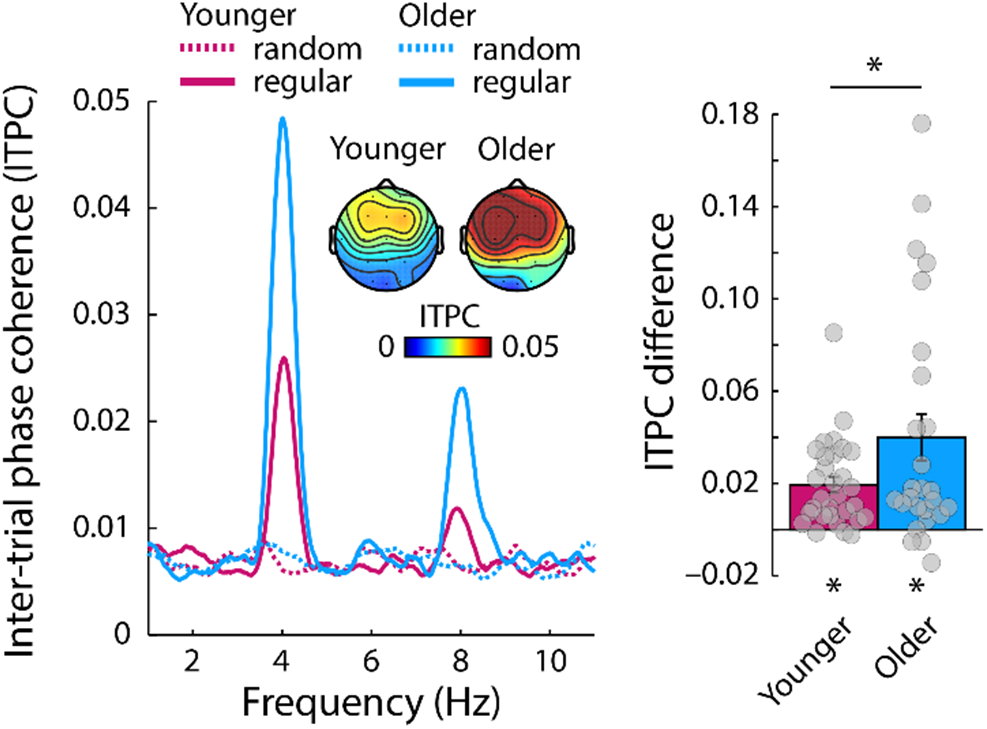
Results for neural synchronization. **Left:** Inter-trial phase coherence (ITPC) as a function of neural frequency for the ‘regular’ and the ‘random’ condition. **Right:** ITPC difference between the ‘regular’ and the ‘random’ condition. Gray dots represent data points for individual participants. *p < 0.05

Notably, the difference in synchronization strength (regular minus random) was particularly prominent for a few older participants (~30%; Figure 5, right). We thus tested whether the distributions’ median differed between age groups. There was no difference between age groups for the median neural synchronization (p = 0.379, r_e_ = 0.120). This suggests that the larger effect of regularity on neural synchronization in older compared to younger people may be due to the neural responses in a subset of older participants.

### Sustained neural activity to temporal regularity

Investigation of the sustained response focused on the time window during which the 4-Hz amplitude modulation could occur in the temporal regularity sounds (1.55–4 s). Figure 6A shows the response time courses and topographies for both conditions (random, regular) and age groups (younger, older). The ANOVA revealed a marginally significant effect of condition (F_1,54_ = 3.340, p = 0.073, η_p_^2^ = 0.058), but no effect of age group (F_1,54_ = 0.325, p = 0.571, η_p_^2^ = 0.006). The condition × age group interaction also tended towards significance (F_1,54_ = 2.838, p = 0.098, η_p_^2^ = 0.050). We had planned tests of simple effects to directly investigate whether low-frequency amplitude modulation in sounds elicits a sustained response in either age group. Accordingly, paired t-tests (contrasting random vs. regular) were calculated separately for younger and older people. The sustained response was larger (i.e., more negative) in the ‘regular’ compared to the ‘random’ condition in younger individuals (t_29_ = 2.335, p = 0.027, r_e_ = 0.398), but not in older individuals (t_25_ = 0.113, p = 0.911, r_e_ = 0.023). As the condition × age group interaction indicates, the response difference between the ‘regular’ and ‘random’ conditions was marginally larger in younger compared to older people (Figure 6B).

**Figure 6:**
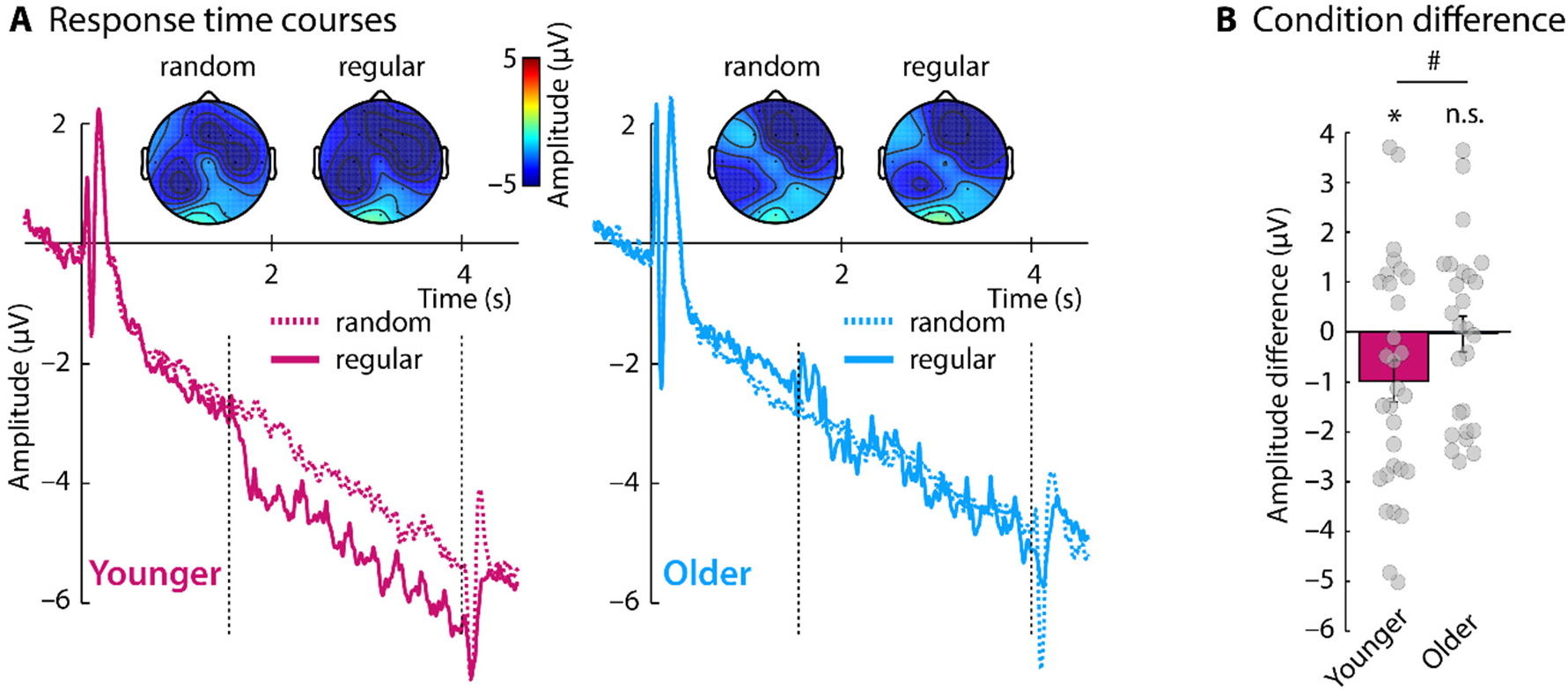
Results for sustained neural activity. **A:** Neural activity time courses for random and regular conditions. Dashed vertical lines mark the time window of interest during which a regular pattern could occur (1.55–4 s) **B:** Difference in sustained response between regular and random conditions. Gray dots represent data points for individual participants. Error bars reflect the standard error of the mean. *p ≤ 0.05, #p ≤ 0.1, n.s. – not significant

### Temporal regularity index

Next, we calculated a regularity index that contrasts the effect of temporal regularity between neural synchronization and sustained neural activity. To this end, we subtracted the response to the ‘random’ condition from the response to the ‘regular’ condition, separately for the sustained response data and the ITPC data. Both differences were separately z-transformed (the sustained response data were also sign-inverted in order to match the sign of ITPC values). The regularity index was then calculated as the subtraction of the z-transformed ITPC effect from the z-transformed sustained response effect. A positive value of this regularity index indicates that the effect of regularity was larger for the sustained response compared to neural synchronization. A negative value indicates that regularity processing was more strongly reflected in neural synchronization compared to sustained activity.

The regularity index was positive and significantly greater than zero for younger individuals (t_29_ = −2.052, p = 0.049, r_e_ = 0.356), but did not differ from zero for older individuals (t_25_ = 1.573, p = 0.128, r_e_ = 0.300). Furthermore, the regularity index differed significantly between age groups. It was significantly more positive for younger compared to older people (t_54_ = 2.505, p = 0.015, r_e_ = 0.323; median test: p = 0.038, r_e_ = 0.279; Figure 7). These results are compatible with the idea that sensitivity to regularity increases from lower-level regions (neural synchronization) to higher-level regions (sustained response) in younger individuals, but that sensitivity to regularity is restricted to lower-level regions in older individuals.

**Figure 7:**
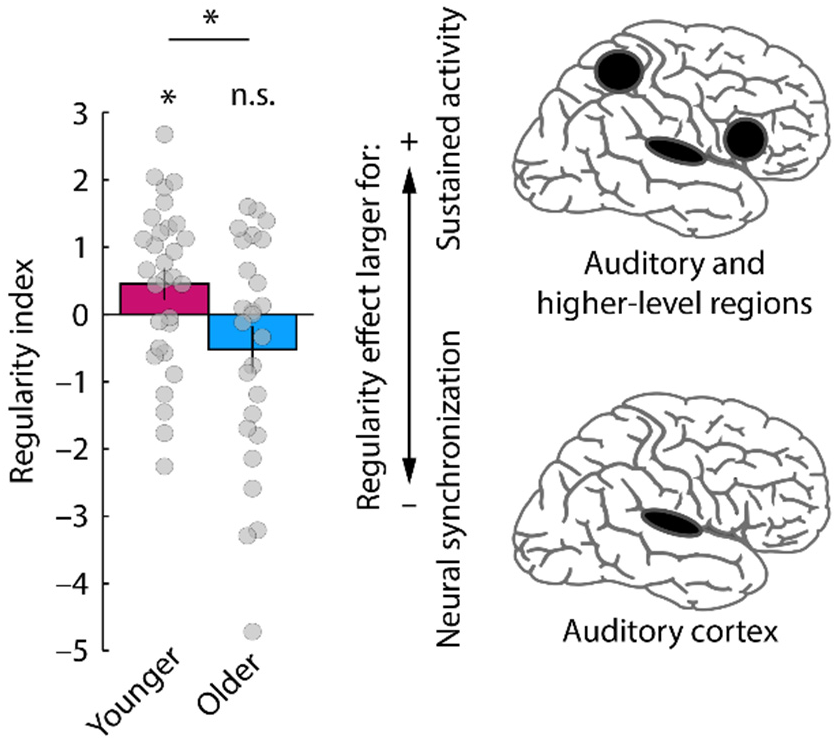
Regularity index reflecting the weighting between the regularity effects for neural synchronization versus the sustained response. Positive values indicate that the effect of regularity was larger for the sustained activity (involving auditory and higher-level brain regions) compared to neural synchronization (involving auditory cortex), and vice versa for negative values. Gray dots represent data points for individual participants. Error bars reflect the standard error of the mean. *p ≤ 0.05, n.s. – not significant

### Correlations between measures of hearing ability and neural responses

We calculated correlations between brain responses and measures of hearing. The following neural measures were used: the linear coefficient from the temporal adaptation paradigm, ITPC regularity effect (regular minus random), regularity effect for sustained activity (regular minus random), and the regularity index. The measures of hearing abilities were as follows: speech in noise score, loudness range (‘very soft’ to ‘very loud’), averaged audiometric threshold for low frequencies (≤2000 Hz). We controlled for multiple comparisons using False Discovery Rate (Benjamini and Hochberg, 1995; Genovese et al., 2002).

None of the correlations between neural measures and measures of hearing were significant when age was partialed out (all p > 0.05). There were also no significant correlations when younger and older participant groups were analyzed separately (all p > 0.05).

## Discussion

In the current study, we investigated age-related changes in central gain and in the neural representations of temporally regular amplitude modulations in sounds. Sensory gain enhancements were assessed by neural response adaptation to sequences of tones in which the onset-to-onset intervals varied. We observed reduced neural adaptation for older compared to younger people, suggesting an increased gain in the aged auditory cortex.

Neural sensitivity to temporal regularity in sounds was assessed by neural synchronization and sustained neural activity to amplitude-modulated narrow-band noises. Neural synchronization to the 4-Hz amplitude modulation of a noise sound – likely reflecting a regularity representation in auditory cortex – was increased in older compared to younger people. In contrast, sustained neural activity – reflecting a regularity representation that likely involves higher-level brain regions in addition to auditory cortex – was diminished in older people. The results may suggest that despite enhanced gain in sensory systems of older individuals, the neural representations of temporal regularity in higher-level brain regions are weakened due to aging and hearing loss.

### Subclinical impairments in hearing accompany aging

In the current study, we assessed the participants’ hearing abilities using audiograms, self-rated reports of speech perception, and measures of loudness perception. Older people had normal audiometric thresholds for low-frequency (≤2000 Hz) pure tones although thresholds were increased by ~9 dB compared to younger people. Sound stimulation in the current study was limited to this low-frequency range and participants had no difficulty perceiving these sounds at supra-threshold levels (Figure 3). However, older people had moderate to severe hearing loss at higher frequencies (≥4000 Hz) and they reported having difficulty understanding speech in the presence of background sound. Hearing abilities were also assessed using a loudness judgment task, during which participants categorized the perceived loudness of pure tones (Epstein and Marozeau, 2006; Al-Salim et al., 2010; Hebert et al., 2013). We also observed loudness recruitment: the loudness range over which older people perceive sounds was reduced compared to younger people (Figure 3; Steinberg and Gardner, 1937; Harris, 1953; Moore et al., 1996; Marozeau and Florentine, 2007).

Elevated hearing thresholds, reduced speech perception in the presence of background sound, and a compressed loudness range suggest subclinical hearing impairments in the population of older people recruited for the current study. Such perceptual changes have been associated with a loss of cochlear compression, that is, with the loss of the active mechanism that in “normal” ears increases sensitivity to soft sounds (Villchur, 1974; Moore and Glasberg, 1993; Moore et al., 1996; Oxenham and Bacon, 2003). The consequences of impaired cochlea function for neural processing upstream in brain stem and cortex can be profound. Changes include a loss of inhibition (Caspary et al., 2008; Takesian et al., 2009, 2012), an increase in excitation (Salvi et al., 2017), and, in turn, an abnormal enhancement of neural responses to sounds (Popelár et al., 1987; Syka et al., 1994; Hughes et al., 2010; Möhrle et al., 2016; Herrmann et al., 2017; Salvi et al., 2017). This and other work therefore suggest that central and peripheral mechanisms are an integrated system, and dysfunction within the system is responsible for loudness recruitment and hyperacusis (Heinz et al., 2005; Cai et al., 2009; Knipper et al., 2013; Zeng, 2013).

### Enhanced neural gain in auditory cortex of older people

In the current study, we measured changes in neural gain between younger and older people by assessing sound-induced neural adaptation in auditory cortex. We utilized a paradigm that allows the assessment of the temporal dynamics of neural adaptation by presenting tone sequences with varying inter-tone intervals (Herrmann et al., 2016). We observed that the response sensitivity to inter-tone interval was larger in older compared to younger people, particularly for longer intervals. The results – similar to those in other reports (de Villers-Sidani et al., 2010; Herrmann et al., 2016; Herrmann et al., 2018) – suggest that, between successive tone presentations, neurons recover faster from adaptation in the aged auditory cortex.

Notably, the level at which sounds were played during EEG recordings was, on average, louder (relative to hearing threshold) for younger compared to older people and slightly above the comfortable loudness level for younger people (Figure 3B). We observed an enhanced neural gain in auditory cortex of older compared to younger people despite the fact that sounds were slightly quieter for older individuals.

The current results are consistent with previous findings in humans and animals that show an age-related enhancement of sound-evoked responses in auditory cortex (Anderer et al., 1996; Tremblay et al., 2003; Sörös et al., 2009; Hughes et al., 2010; Bidelman et al., 2014; Herrmann et al., 2018). Our results are also in line with the observation that damage to the auditory periphery in animals leads to hyperresponsivity in the afferent auditory system (Gerken, 1979; Popelár et al., 1987; Syka et al., 1994; Möhrle et al., 2016; Salvi et al., 2017). In fact, denervation of any biological structure appears to lead to hyperresponsivity (Cannon and Rosenblueth, 1949; Larrabee, 1949; Gerken, 1979) and the observed effects may reflect a general property of denervated tissue. For the auditory sensory modality, gain enhancements in mammalian brain structures may be the result of a loss of inhibition (Caspary et al., 2008; Takesian et al., 2009; Llano et al., 2012; Takesian et al., 2012; Auerbach et al., 2014) and an increase in excitation (Salvi et al., 2017).

An enhanced gain in older people may have a variety of consequences. Firstly, neuronal firing is metabolically expensive (Attwell and Laughlin, 2001) and an enhanced gain may thus lead to increased energy consumption. In addition, large neural responses are thought to reflect uncertainty about the world compared to small responses, which are thought to reflect response cancellation due to internally generated predictions (Friston, 2005; Friston and Kiebel, 2009). According to this framework, a gain enhancement in older people would indicate that the aged auditory cortex responds stereotypical and abnormally rigorous to external events rather than being weighted by internally generated predictions. Furthermore, an increase in neural gain appears to manifest as altered neural adaptation to sounds (Herrmann et al., 2016) and sound statistics (Herrmann et al., 2018), which may impair the flexible adjustment of perceptual systems to different acoustic environments, and may underlie the challenges older people experience with filtering out irrelevant information. Finally, an increase in neural gain in auditory cortex has been related to impaired speech perception (Millman et al., 2017; Goossens et al., 2018). Hence, despite older people showing enhanced responsivity to sound in auditory cortex, acoustic environments may not be represented accurately in the brain.

### Neural sensitivity to temporal regularity in aging

We investigated the effects of age on two different neural signatures of the processing of temporal regularity in sounds – neural synchronization and sustained neural activity. We observed that neural synchronization was enhanced in about 30% of older individuals. This observation is consistent with enhanced neural synchronization observed previously for sounds that contain low-frequency amplitude modulations (Purcell et al., 2004; Goossens et al., 2016; Presacco et al., 2016a, b; Herrmann et al., 2017; Lai et al., 2017). The fact that the current effects appear somewhat weaker compared to previous work may be due to the slightly perceptually quieter level used for older compared to younger adults (previous work either used louder sounds for older people or equally loud sounds). Moreover, data points for individual participants are commonly not shown, which precludes assessment of individual differences in synchronization magnitude in the older group.

Previous source localizations suggest that synchronization of neural activity with low-frequency temporal regularities in sounds is strongest in auditory cortex (Herrmann et al., 2013a; Keitel et al., 2017; Millman et al., 2017). The finding of enhanced synchronization in auditory cortex for older people is consistent with the age-related gain enhancement that we observed using the adaptation paradigm. Furthermore, modeling work suggests that changes in neural synchronization may be due to changes in adaptation properties of neurons (Ladenbauer et al., 2012; Augustin et al., 2013), but this is a topic that requires more research.

In contrast to neural synchronization, stronger sustained neural activity in response to the sound’s amplitude modulation was present in younger individuals, but absent in older people (with the difference between age groups approaching significance; Figure 6). Topographical distributions of the sustained activity were widespread and may suggest the involvement of higher-level brain regions: this is consistent with previous source localizations showing that hippocampus, frontal cortex, and parietal cortex are part of the network underlying regularity-related sustained activity (Tiitinen et al., 2012; Barascud et al., 2016; Teki et al., 2016) in addition to auditory cortex (Pantev et al., 1994; Pantev et al., 1996; Gutschalk et al., 2002). Our data may therefore suggest that the processing of temporal regularities in a network involving higher-level brain regions is altered in older people. The regularity index, which reflects the relative weight of sustained neural activity versus neural synchronization for the processing of temporal regularity (Figure 7) suggests that aging (and hearing loss) is associated with an over-representation of temporal regularity in auditory cortex, and with an under-representation in the network involving higher-level brain regions.

The current data indicate that a reliable sensory representation may not be sufficient for the processing of temporal regularity. Although an increase in neural gain – which is thought to compensate for degraded inputs from the auditory periphery – may support the detection of weak signals (Gerken, 1979; Ernst and Moore, 2012; Chambers et al., 2016; Schlittenlacher and Moore, 2016), it may affect how a sound’s regularity is represented in cortical structures, in an unhelpful way. This is in line with previous work demonstrating a correlation – with age partialed out – between neural gain enhancements and decreased speech perception performance in the presence of modulated background sound (Millman et al., 2017; Goossens et al., 2018), and with the observation that hearing loss increases the perceived magnitude of low-frequency amplitude modulation in sounds (Moore et al., 1996). This previous work together with the current findings indicate that neural function in central auditory brain structures of older people appears fundamentally altered, including, among others, a change in sensitivity to a sound’s temporal regularity in auditory cortex and possibly in higher-level brain regions.

## Conclusions

In the current study, we investigated how aging affects the sensitivity of cortical structures to temporal regularity – here, amplitude modulation – in sounds. Synchronization of auditory cortex activity with the amplitude modulation of a narrow-band noise was enhanced in about 30% of older compared to younger people. Despite the reliable representation of temporal regularity in auditory cortex (as indicated by neural synchronization), sustained neural activity – a neural signature of regularity processing that probably involves higher-order brain regions such as frontal and parietal cortex – was diminished in older people. The data suggest that aging leads to an over-representation of low-frequency amplitude modulation in auditory cortex at the cost of a diminished representations of amplitude modulation in higher-order cortical areas.

## Acknowledgements

Research was supported by the Canadian Institutes of Health Research (MOP133450 to I.S. Johnsrude). BH was supported by a BrainsCAN postdoctoral fellowship (Canada First Research Excellence Fund; CFREF).

